# Genome-wide association analysis with a 50K transcribed gene SNP-chip identifies QTL affecting muscle yield in rainbow trout

**DOI:** 10.1101/355792

**Authors:** Mohamed Salem, Rafet Al-Tobasei, Ali Ali, Daniela Lourenco, Guangtu Gao, Yniv Palti, Brett Kenney, Timothy D. Leeds

## Abstract

Detection of coding/functional SNPs that change the biological function of a gene may lead to identification of putative causative alleles within QTL regions and discovery of genetic markers with large effects on phenotypes. Two bioinformatics pipelines, GATK and SAMtools, were used to identify ~21K transcribed SNPs with allelic imbalances associated with important aquaculture production traits including body weight, muscle yield, muscle fat content, shear force, and whiteness in addition to resistance/susceptibility to bacterial cold-water disease (BCWD). SNPs were identified from pooled RNA-Seq data collected from ~620 fish, representing 98 families from growth- and 54 families from BCWD-selected lines with divergent phenotypes. In addition, ~29K transcribed SNPs without allelic-imbalances were strategically added to build a 50K Affymetrix SNP-chip. SNPs selected included two SNPs per gene from 14K genes and ~5K non-synonymous SNPs. The SNP-chip was used to genotype 1728 fish. The average SNP calling-rate for samples passing quality control (QC; 1,641 fish) was ≥ 98.5%. Genome-wide association (GWA) study on 878 fish (representing 197 families from 2 consecutive generations) with muscle yield phenotypes and genotyped for 35K polymorphic markers (passing QC) identified several QTL regions explaining together up to 28.40% of the additive genetic variance for muscle yield in this rainbow trout population. The most significant QTLs were on chromosomes 14 and 16 with 12.71% and 10.49% of the genetic variance, respectively. Many of the annotated genes in the QTL regions were previously reported as important regulators of muscle development and cell signaling. No major QTLs were identified in a previous GWA study using a 57K genomic SNP chip on the same fish population. These results indicate improved detection power of the transcribed gene SNP-chip in the target trait and population, allowing identification of large-effect QTLs for important traits in rainbow trout.

## Introduction

Aquaculture provides sustainable production of food fish with high protein/low-saturated fat to satisfy increasing U.S. and worldwide demand. To enable increased production by the aquaculture industry and to meet the ever-growing demand for fish, we need fast/efficient growth and high-quality fillets. However, a major constraint to increasing production efficiency is the lack of genetically improved strains of fish for aquaculture [1; 2]. Development of tools that will enable genomic selection for improved aquaculture production traits will greatly benefit the aquaculture industry.

Fast/efficient muscle growth is a major trait affecting profitability of the aquatic muscle food industry. The genetic basis of muscle growth traits is not well studied in fish. Understanding molecular mechanisms of fish muscle growth can facilitate broodstock selection decisions. In addition, fish fillets are nutritionally and economically valuable products. Skeletal muscle is the most abundant tissue and edible portion of fish and typically constitutes about 50-60% of the fish weight [3]. Growth, development and quality traits of muscle are governed by organized expression of genes encoding contractile and regulatory proteins [4].

Genomic selection (GS) tools have been developed to increase the efficiency of genetic improvement in livestock compared to conventional pedigree-based selective breeding methods[5]. This concept has been recently demonstrated for bacterial cold-water disease (BCWD) resistance in rainbow trout aquaculture [6]. Marker-assisted selection (MAS) can be used to improve breeding for phenotypes with large-effect QTLs. This method has been recently applied for the trait of infectious pancreatic necrosis virus (IPNV) resistance in Atlantic salmon [7]. Genetic maps, characterizing the inheritance patterns of traits, and markers have been developed and used for a wide range of species, including fish. These tools target the discovery of allelic variation affecting traits with an ultimate goal of identifying DNA sequences underlying phenotypes [8]. Markers have been identified with a variety of molecular techniques. Single nucleotide polymorphisms (SNPs) are abundant and distributed genome-wide, therefore, they are most suitable for high-throughput association studies [9; 10]. SNPs located within or near coding sequences, cSNPs, are especially important because they have the potential to change protein function [11; 12; 13]. Therefore, cSNPs are particularly useful as genetic markers with large-effect on phenotypes, allowing MAS and improved accuracy of whole-genome selection. Because the muscle yield trait targeted in this study requires lethal sampling to measure the phenotype, only family-specific EBVs are available for breeding candidates in traditional breeding programs. The ability to use genomic selection or MAS will allow further within-family selection for the muscle yield trait, and thus is anticipated to increase the accuracy of genetic predictions and selection response.

Recently, we used an RNA-Seq approach to identify putative SNPs with allelic imbalances associated with total body weight, muscle yield, muscle fat content, shear force, and whiteness [12; 13]. Similarly, RNA-Seq data were used to identify SNPs with allelic imbalances in fish families showing variations in resistance to *Flavobacterium psychrophilum*, the etiological agent of BCWD in rainbow trout [14; 15]. Together about 50K and 229K transcribed SNPs were identified in the two studies, respectively. Of them, ~21K SNPs had allelic-imbalances in families with contrasting phenotypes. The first objective of this study was to design, develop, and validate a 50K transcribed gene SNP-chip. The chip content includes the 21K transcribed SNPs with allelic-imbalances associated with the aforementioned traits and ~29K SNPs without allelic-imbalances that were strategically added to achieve more even genome-wide distribution. The new SNP-Chip is available from Affymetrix. The second objective of this study was to test the feasibility of using the new SNP-chip in GWA analysis to identify QTL explaining muscle yield variance in the USDA/NCCCWA rainbow trout growth-selected line. The results were compared with a previous GWA study for the same trait in the same population that we have previously conducted with a genomic-based 57K SNP chip [10].

## Materials and methods

### ETHICS STATEMENT

Institutional Animal Care and Use Committee of the United States Department of Agriculture, National Center for Cool and Cold Water Aquaculture (Leetown, WV) specifically reviewed and approved all husbandry practices and experimental procedures used in this study (Protocols #056 and 076).

### SOURCE AND SELECTION OF SNPS FOR THE CHIP

Recently, we used RNA-Seq and two bioinformatic pipelines, GATK and SAMtools, for discovering coding/functional SNPs from 98 rainbow trout fish families (5 fish each) showing variations in whole-body weight, muscle yield, muscle fat content, shear force, and whiteness [13]. GATK detected 59,112 putative SNPs and SAMtools detected 87,066 putative SNPs. The two datasets contained approximately 50K non-redundant common SNPs; of which, 30,529 mapped to protein-coding genes (with 7.7% non-synonymous SNPs) and 4,386 mapped to lncRNAs. A total of 7,930 non-redundant SNPs had allelic imbalances between the low- and high-ranked families for the phenotypes. Validation of a subset of 92 SNPs revealed 1) 86.7-93.8% success rate in identifying polymorphic SNPs and 2) 95.4% consistent matching between DNA and cDNA genotypes, indicating a high rate of identifying SNPs using RNA-Seq. This SNP data set was recently published and is available through the NCBI dbSNP database (accession numbers ss#2711191806-2711287038 in addition to ss#2137497773) [13].

Similarly, we identified transcribed gene SNPs in two genetic lines, ARS-Fp-R (resistant) and ARS-FP-S (susceptible), that were created by selective breeding to exhibit divergent resistance to BCWD. RNA-Seq analysis of pooled RNA samples was used to identify SNPs from the resistant and susceptible genetic lines. Fish belonging to resistant and susceptible genetic lines were collected on day 1 and day 5 post-challenge with Fp versus PBS injection [14; 15]. Using GATK bioinformatics pipelines, ~229K transcribed SNPs were identified[14]. The total number of SNPs with allelic imbalance, after removing redundant SNPs, was 7,951.

The SNPs identified in the previous two studies were used as a source to build the SNP array described in this study. About 21K transcribed SNPs with allelic-imbalances associated with the above-listed traits were included in the chip. These SNPs were identified from pooled RNA-Seq data collected from ~620 fish, representing 98 families from the ARS growth-selected line and 54 families from the ARS-Fp-R and -S lines. In addition, about 29K transcribed SNPs without allelic-imbalances were selected from all the putative SNPs and were strategically added to the chip with the aim of achieving even distribution of SNPs along the rainbow trout 29 chromosomes. The additional SNPs were selected to represent as many genes as possible in the genome: two SNPs were selected per gene from 14K genes with available SNPs. The chip includes ~5K non-synonymous SNPs. The chip has probe sets for a total number of 50,006 SNPs.

### CHIP GENOTYPING QUALITY ASSESSMENT

The SNP-chip was used in genotyping 1,728 fish from the USDA-ARS genetic lines. The Affymetrix SNPolisher software was used to calculate the chip SNP- and sample-metrics and assess QCs and filter samples/genotypes at the default setting [16]. Forty-seven SNPs previously genotyped by a Fluidigm PCR-based assay [13] were used to check quality of Affymetrix chip genotyping using 120 samples genotyped by both the chip and Fluidigm SNP assays. In addition, we confirmed the quality of the SNPs and the order of the samples included in the genotyping panel through pedigree check. Among the fish genotyped we included previously confirmed parental-pairs of nine families with 470 offspring and confirmed an average of 99.4% matching between offspring SNP genotypes and the genotypes of the expected parents.

### SNP GENOMIC DISTRIBUTION AND ANNOTATION

SNPs used in building the chip were identified using the first draft of the rainbow trout reference genome [14]. To update genomic coordinates according to the newly released genome assembly (GenBank assembly Accession GCA_002163495, RefSeq assembly accession GCF_002163495) [17], SNPs were mapped by BLASTing the SNP probe sequences (70 nt) to the new genome sequence. Sequences with 100% identity match and no gap with single hits were assigned to the new genome position. Sequences with multiple hits were re-Blasted using probe size of 150 nt by adding 40 nt flanking sequence in both direction. A total of 45K SNPs out of 50K SNPs were successfully assigned to the new genome and were used for the GWA analyses.

SNPeff program was used to classify and annotate functional effects of the SNPs [18]. The gff file of the new rainbow trout genome reference was used to determine position of the SNPs in a gene i.e. located within mRNA start and end positions (genic), within a CDS, 5’UTR or 3’UTR. SNPs not within start and end positions of mRNA were considered intergenic. Upstream/ downstream intergenic SNPs were determined if located within 5 Kb of an mRNA. SNPs within lncRNAs were determined using gtf file of our previously reported lncRNA reference [63]. SNP annotation was performed by intersecting the SNPs bed file with the gff/gtf file using Bedtools software [19].

### RAINBOW TROUT POPULATION AND PHENOTYPES USED FOR GWA

Genome-wide association analysis was carried out using fish from a growth-selected line that has been previously described [20]. Briefly, this synthetic line is a 2-yr-old winter/spring-spawning population that was developed beginning in 2002, became a closed population in 2004, and since then has gone through 5 generations of genetic selection for improved growth performance. Fish from two consecutive generations (i.e., the third and fourth generations of growth selection) were included in this study. Phenotypic data and DNA samples were collected from 878 fish (representing 98 families from year-class (YC) 2010 and 99 families from YC 2012). Methods used to sample fish from each nucleus family and to characterize muscle yield have been described previously [10]. Eggs were hatched in spring water at 7-13°C to synchronize hatch times. Each family was stocked separately in 200-L tanks and hand-fed a commercial fishmeal-based diet beginning at swim-up. Neomales were developed from a subset of alevins from the previous year class by feeding 2 mg/kg of 17α-methyltestosterone for 60 d post-swim-up, and the masculinized females were used as sires for the following generation. At 5-months old, fish were uniquely tagged by inserting a passive integrated transponder, and tagged fish were combined and reared in 1,000-L communal tanks. Fish were fed a commercial fishmeal-based diet using automatic feeders. EBV were computed based on a two-trait model, 10-mo BW and thermal growth coefficient (TGC), using MTDFREML [21]. Each generation, EBV was used as selection criterion and mating decisions were made to maximize genetic gain while constraining the inbreeding rate to ≤1% per generation using EVA evolutionary algorithm [22]. Data from masculinized fish were not used in the growth analysis.

Fish were harvested between 410 and 437 days post-hatch (mean body weight = 985 g; SD = 239 g), between 446 and 481 days post-hatch (mean body weight = 1803 g; SD = 305 g), for the 2010, and 2012 hatch years, respectively. Individual body weight data were recorded at harvesting. Fish were taken off feed 5 days before harvesting. For measurement of muscle yield when harvested at each of five consecutive weeks, approximately 100 fish (i.e., 1 fish per full-sib family per week) were anesthetized in approximately 100 mg/L of tricaine methane sulfonate (Tricaine-S, Western Chemical, Ferndale, WA) slaughtered, and eviscerated. Head-on gutted carcasses were packed in ice, transported to the West Virginia University Muscle Foods Processing Laboratory (Morgantown, WV), and stored overnight. The next day, carcasses were manually processed into trimmed, skinless fillets by a trained faculty member and weighed; muscle yield was calculated as a percent of total body weight [23].

### GWA ANALYSES

Weighted single-step GBLUP (WssGBLUP) was used to perform GWA analysis as implemented in previous studies [24; 25; 26]. In addition to phenotypic data, wssGBLUP integrates genotype and pedigree information to increase estimation precision and detection power [25] in a combined analysis that is executed by the BLUPF90 software [27].

The following mixed model was used for single trait analysis:

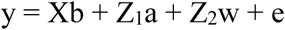

where y is the vector of the phenotypes, b is the vector of fixed effects including harvest group and hatch year, a is the vector of additive direct genetic effects (i.e., animal effect), w is the vector of random family effect, and e is the residual error. The matrices X, Z_1_, and Z_2_are incidence matrices for the effects contained in b, a, and w, respectively.

As BLUP considers the variance components are known, AIREMLF90 [27] was used to estimate variance components for the additive direct genetic effect, random family effect, and residuals. Inbreeding was considered in all analyses, and was calculated using INBUPGF90 [27] on 63,808 fish that represent five generations in the NCCCWA population. Quality control (QC) of genomic data was performed using BLUBF90 [27] with the following parameters: SNP with minimum Allele Frequency (MAF) >0.05, SNP with call rate > 0.90, animals with call rate >0.90, and SNP with a difference between observed and expected allele frequency <0.15 (i.e., HWE test) were kept in the data. Out of a total of 50,006 SNPs, 35,322 SNPs passed QC. For the first iteration of WssGBLUP, all SNPs were assigned the same weight (e.g., 1.0). For the next iteration, weights were calculated based on the SNP effects (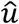) estimated in the previous iteration as 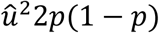, where p is the current allele frequency. Each iteration was performed using three steps as follows: first, weight was assigned as described above; second, BLUPF90 [27] was used to compute genomic estimated breeding values (GEBV) based on a realized relationship matrix (**H**) that combines pedigree (**A**) and genomic relationship matrix (**G**), the last considered weights for SNP; and third, postGSF90 [27] was used to calculate SNP effects and weights based on sliding windows of 50 adjacent SNPs. A total of 2 iterations were used. A window based on physical size (e.g. 1Mb) was not used to avoid biases due to uneven distributed SNPs in the new SNP chip. A Manhattan plot based on the proportion of additive genetic variance explained by the windows was created using the qqman package in R [28]; thus, the genomic windows explaining significant proportion of the additive genetic variance for muscle yield could be detected.

### CITRATE SYNTHASE (CS) ACTIVITY ASSAY

GWA analysis (described below) showed a SNP window contained the CS gene associated with the genetic variance in muscle yield. To assess the potential effect of the SNPs in this gene, we measured the CS activity in 100 fish from the 2012 year-class. Frozen muscle tissue samples were homogenized using electric homogenizer on ice followed by centrifugation at 1,000g for 15 min at 4 °C. The supernatant was used to assess the total protein concentration and CS activity. Total protein concentration was assessed using a BCA protein assay kit at 562 nm with bovine serum albumin (BSA) as the standard. CS activity was determined from the rate of appearance of reduced DTNB (5,5′-dithiobis [2-nitrobenzoic]), which was monitored with a spectrophotometer at 412 nm[29]. For the CS assay, 10 μL of diluted tissue homogenate (1.0 mg/ ml) was incubated with 140 μL reaction medium (0.1mM DTNB, 0.2mM AcetylCoA, 0.15mM oxaloacetic acid, pH 8.0). The absorbance was read in triplicate at 412 nm (25 °C) after 4 min. CS activity was expressed as ∆OD/ mg protein.

## Results and discussion

### CHIP GENOTYPING QUALITY ASSESSMENT

The SNP-chip was used to genotype 1,728 fish. Out of 50,006 SNPs, 32,273 SNPs (64.5%) were characterized as high quality and polymorphic and 3,458 SNPs (6.9%) were high quality monomorphic (Table 1).

**Table 1.**
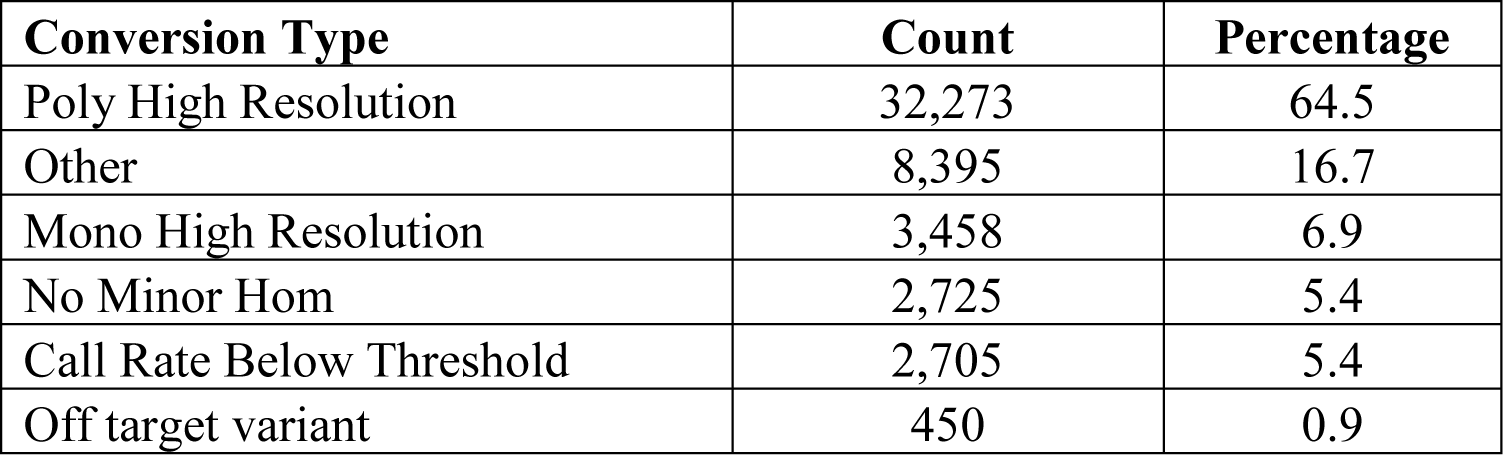
SNP chip Metric summary

| Conversion Type | Count | Percentage |
| --- | --- | --- |
| Poly High Resolution | 32,273 | 64.5 |
| Other | 8,395 | 16.7 |
| Mono High Resolution | 3,458 | 6.9 |
| No Minor Hom | 2,725 | 5.4 |
| Call Rate Below Threshold | 2,705 | 5.4 |
| Off target variant | 450 | 0.9 |

The Affymetrix SNPolisher software was used to filter samples/genotypes at the default setting [16]. Out of 1,728 genotyped samples, 1,641 (94.9%) fish samples were retained, and 87 samples were filtered out because they failed to meet the 0.97 call rate (CR) and 0.82 Dish QC (DQC) thresholds. The average QC call rate for the passing samples was 99.6% (Table 2).

**Table 2.** SNP chip Sample QC Summary

|  |  |
| --- | --- |
| Number of input samples | 1,728 |
| Samples passing DQC | 1,722 |
| Samples passing DQC and QC CR | 1,641 |
| Samples passing DQC, QC CR and Plate QC | 1,641 (94.9%) |
| Number of failing samples | 87 |
| Number of Samples Genotyped | 1,641 |
| Average QC CR for the passing samples | 99.66 |

We compared the Affymetrix genotyping results of 47 SNPs that were previously genotyped by a Fluidigm PCR-based assay [13]. Using 120 samples genotyped by both methods, there was a 99.5% match in genotypes between the two assays for high-resolution polymorphic markers (data not shown). This test demonstrates the high quality of the SNP chip and reliable genotyping data for the subsequent GWA analyses.

The SNP-chip showed an average minor allele frequency (MAF) of 0.25 and standard deviation of 0.134. A total of 27,280 SNPs had MAF> 0.1 and 16,101 SNP more than 0.25 (Figure 1).

**Figure 1.**
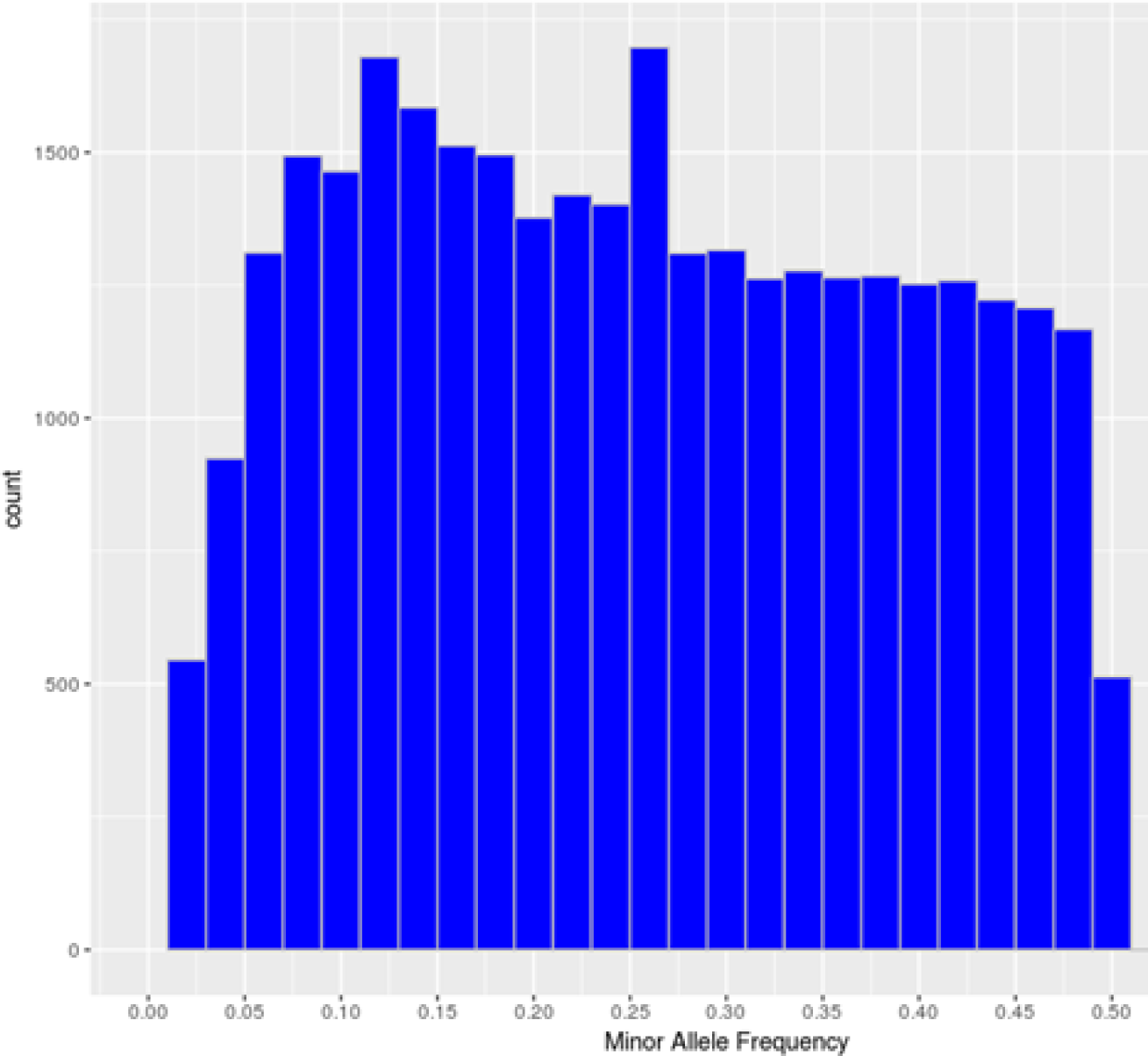
Minor allele frequency distribution of the polymorphic high-resolution SNPs in the SNP chip.

### SNP DENSITY AND GENOMIC DISTRIBUTION

SNPs used in this study to build the chip were initially identified using a rainbow trout reference genome published by Berthelot et al. in 2014 [30]. However, in this reference only ~1 Gb out of a 2.1 Gb total length of the assembly is anchored to chromosomes. Recently, a newer genome assembly has been built that is currently available at NCBI (Accession GCA_002163495) [17]. The new assembly has a 1.94 Gb total length (89% of the genome) anchored to 29 chromosomes. A total of 45,590 SNPs out of 50,006 existing in the SNP-chip were mapped to the new genome assembly with an average of 1,572 SNPs per chromosome. The average SNP density was 1 SNP per 42.7 Kb, with a range of 1 SNP/33.5 Kb (Chromosome 16) to 1 SNP/61.6 Kb (Chromosome 23). Figure 2 shows the number of SNPs per chromosome and the SNP density distribution. A total of 21K out 50K SNPs on the chip were selected based on putative association with phenotypic traits, and hence, were expected to be clustered in specific genome loci. However, supplementing the chip with 29K SNPs (two SNPs per gene) perhaps helped in randomizing the SNP distribution in the genome. Previously, a 57K genome-wide SNP array for rainbow trout reported an average of 1,551.4 mapped SNPs per chromosome [10]. The 57K array was designed primarily using SNPs originating from RAD-Seq sequencing of doubled-haploid clonal lines [31] and whole genome re-sequencing of fish from the Aquagen (Norway) breeding program. A key point here, is that the

SNPs included in the 57K chip were originated from other genetic lines. Hence, although polymorphic enough in the NCCCWA growth line used in this study for conducting GWA as we have previously shown[10], the SNPs used for GWA in this study were originated from the investigated population and were expected to be more informative due to ascertainment bias [32].

**Figure 2.**
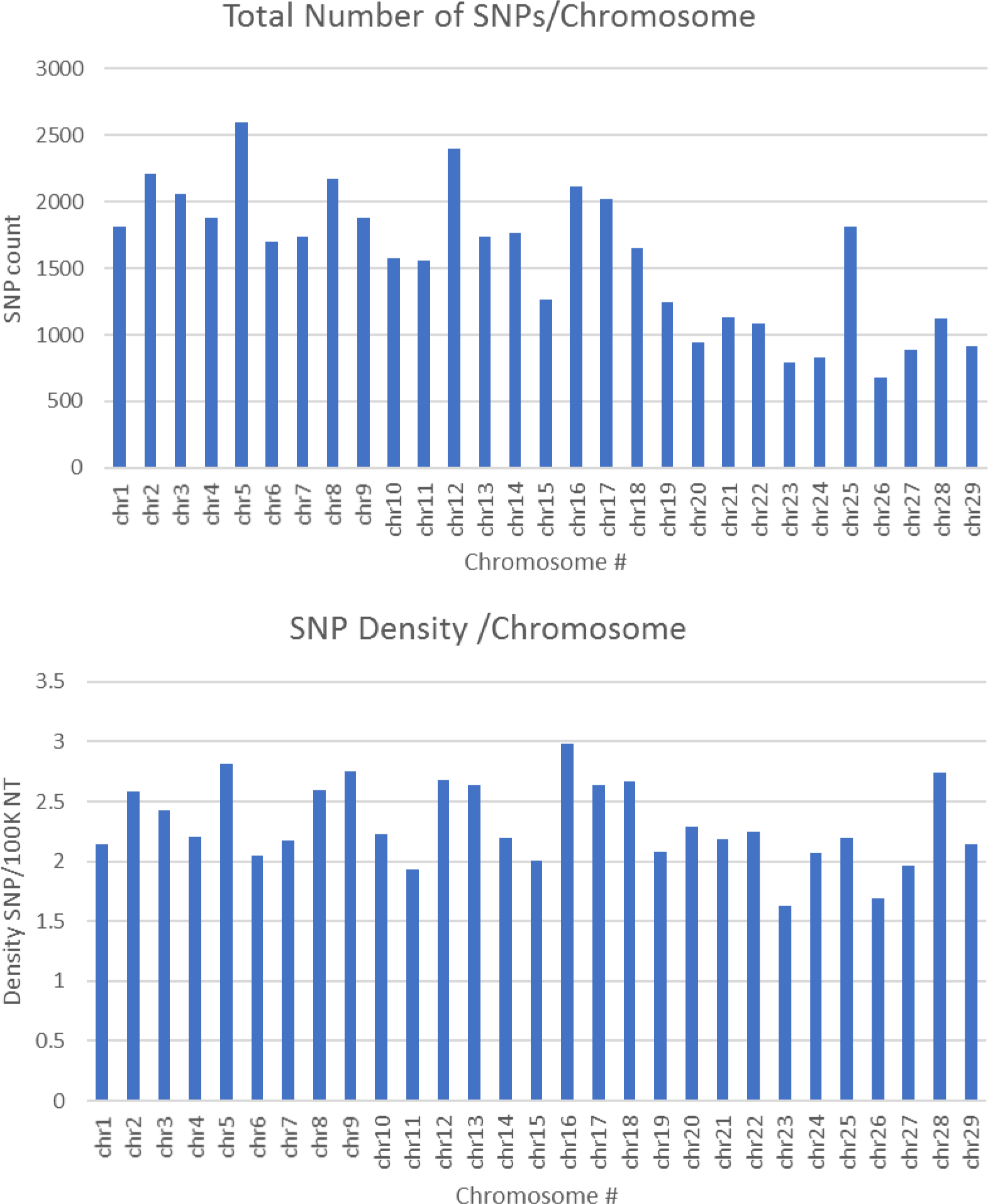
Number of SNPs per chromosome and SNP density distribution (SNP/100K nucleotide).

### SNP ANNOTATION AND CLASSIFICATION BASED ON FUNCTIONAL EFFECTS

SNPeff program was used to classify and annotate functional effects of the SNPs. A total of 45,590 SNPs were included in this analysis. Classifying SNPs by impact showed 636 effects (0.23%) with high impact (stop-gain) and 20,987 effects (7.86%) with moderate impact, (missense variants). The rest (91.9%) represents low to moderate variant effect including synonymous and non-coding SNPs. Figure 3 shows percent of SNP effects by gene regions. A total of 32.8% of the effects were within transcripts with 16.5% exonic, 1.3% in the 5’-UTR and 12.8% in 3’-UTR. All SNPs on the chip were identified through transcriptome sequencing. Surprisingly, there were 14% upstream and 18.1% downstream effects (within 5 Kb of the genes). The upstream/downstream percent is consistent with our previous report that showed 17.1-20.2% SNPs within 5 Kb upstream/downstream of protein-coding genes in one of two SNP data sets used in building the SNP-chip [13]. On the other hand, there was only 1.9% of the SNP effects within intergenic regions, compared to 37.7-49.2% intergenic SNPs in the previous study[13]. In our previous study, the high percentage of intergenic and upstream/downstream SNPs was explained by the incomplete annotation of protein-coding genes and exons used in the previous version of the rainbow trout reference genome [30]. The drop in the percentage of intergenic SNP effects in this study may be due to the improved gene annotation of the current version of the genome reference.

**Figure 3.**
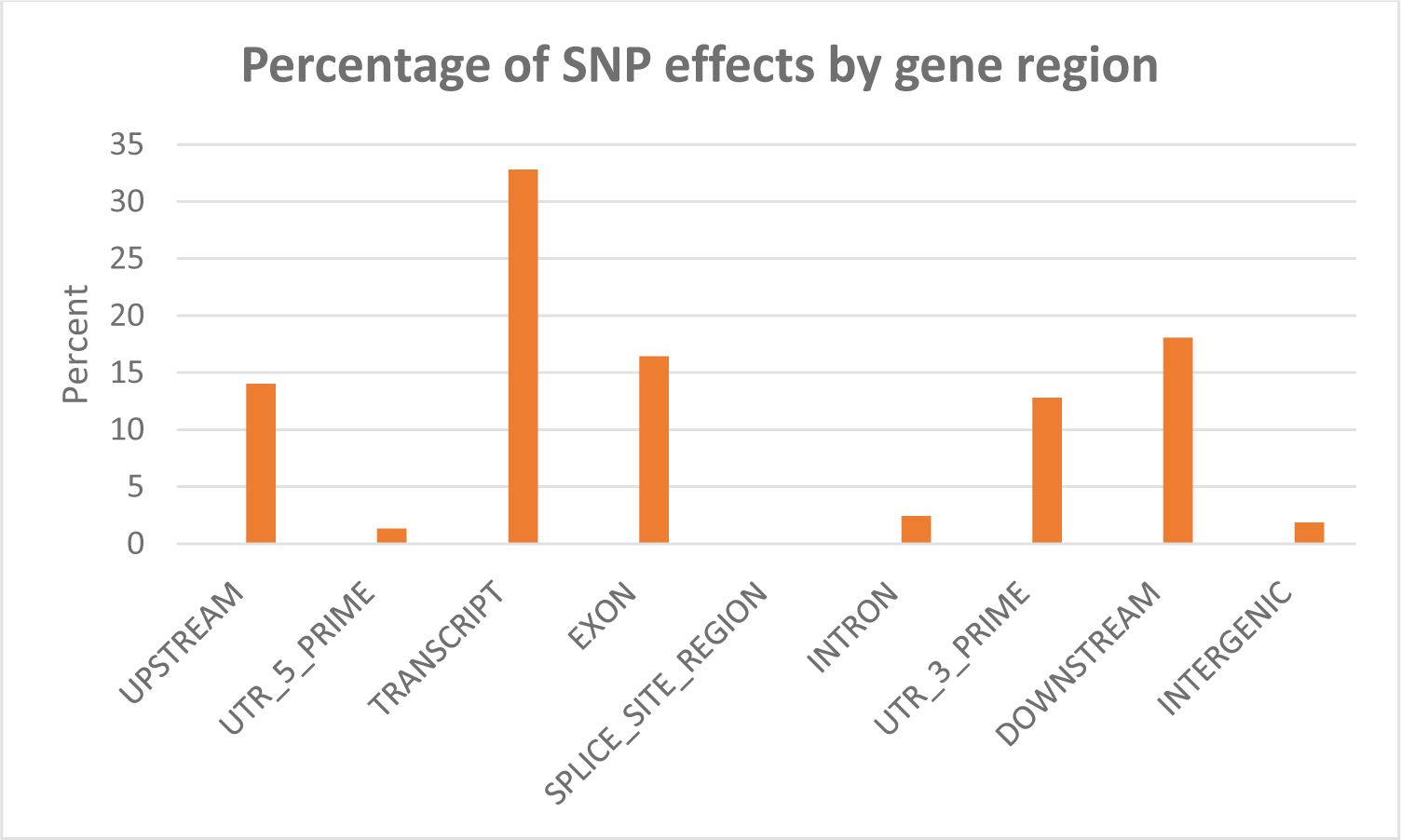
Percentage of SNP effects by gene region.

### GWA ANALYSES

#### Genomic regions associated with muscle yield

GWA analysis using WssGBLUP identified 163 SNPs, each explaining at least 2% of the genetic variance of muscle yield (Tables 3 and 4; Supplementary data sheet 1). The SNPs were clustered into 4 main chromosomes (14, 16, 9, and 17). Chromosomes 14 and 16 showed the highest peaks with genomic loci explaining up to 12.71% and 10.49% of the genetic variance, respectively. The total variance explained by these loci is 23.2%. Figure 4 shows a Manhattan plot displaying association between SNP genomic sliding window of 50 SNPs and muscle yield. Sixty-nine of the 163 SNPs (42.2%) were previously identified as SNPs with allelic imbalances associated with muscle yield in the original SNP data set used to build the SNP chip [13]. Twenty-one of the 163 SNPs caused nonsynonymous mutations. The rest of the SNPs were either silent mutations or located in UTR of the genes indicating their potential epigenetic mechanism of gene regulation. Important SNPs with more than 5% genetic variance are discussed below and all 163 SNPs are listed in Supplementary data sheet 1.

**Table 3.**
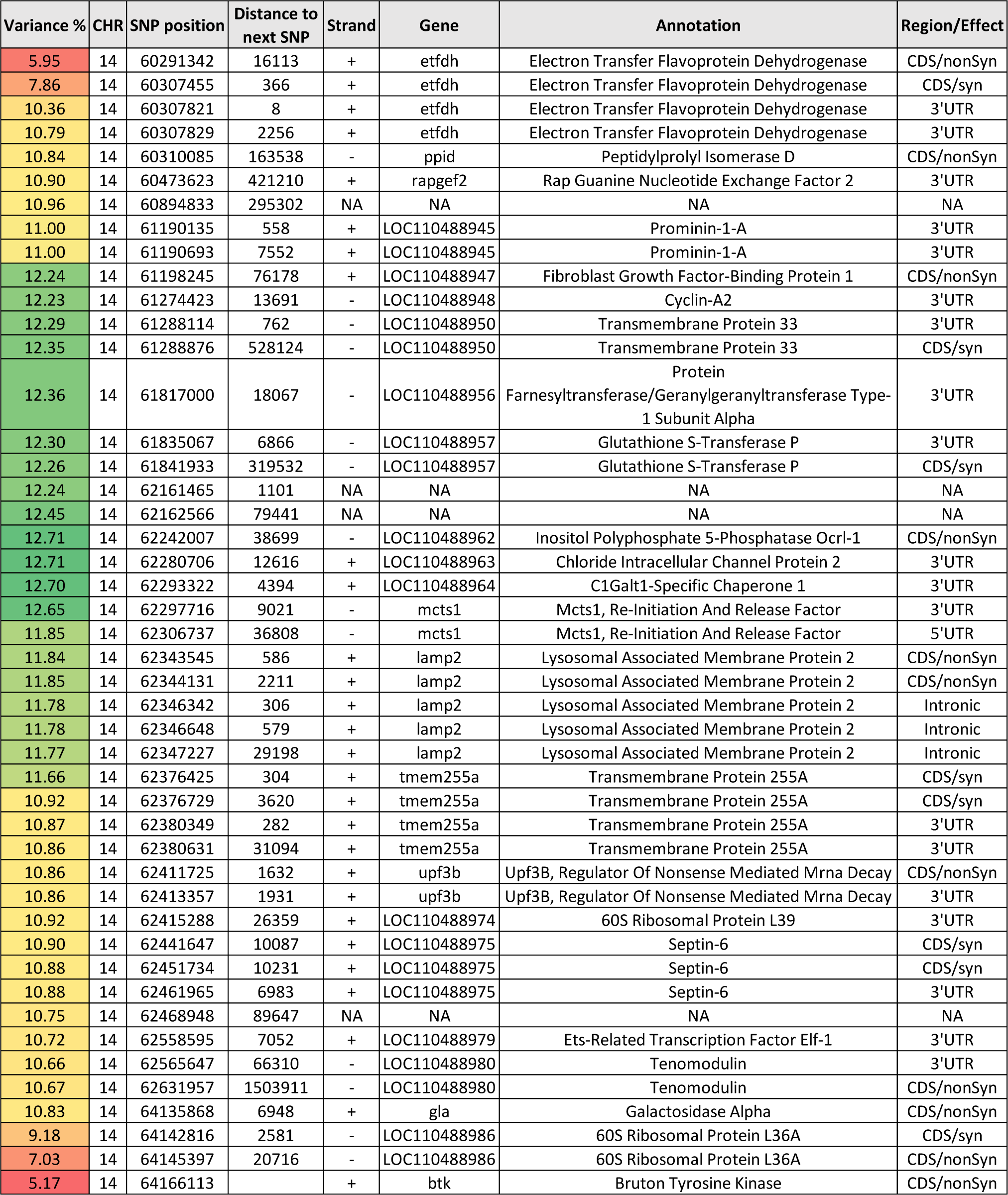
Selected SNP markers explaining the largest proportion of genetic variance (>5%) for muscle yield in chromosome 14 using 50 adjacent SNP windows. Color intensities reflect changes in additive genetic variance (green is the highest and red is the lowest).

**Table 4.** Selected SNP markers explaining the largest proportion of genetic variance (>5%) for muscle yield in chromosome 16 using 50 adjacent SNP windows. Color intensities reflect changes in additive genetic variance (green is the highest and red is the lowest).

| Variance % | CHR | SNP position | Distance to next SNP | Strand | Gene | Annotation | Region/Effect |
| --- | --- | --- | --- | --- | --- | --- | --- |
| 4.62 | 16 | 39953311 | 12000 | + | tnfrsf5a | Tnf Receptor Superfamily Member 5A Precursor | 5'UTR |
| 5.09 | 16 | 39965311 | 3 | + | tnfrsf5a | Tnf Receptor Superfamily Member 5A Precursor | CDS/nonSyn |
| 6.03 | 16 | 39965314 | 689 | + | tnfrsf5a | Tnf Receptor Superfamily Member 5A Precursor | CDS/nonSyn |
| 6.83 | 16 | 39966003 | 608 | + | tnfrsf5a | Tnf Receptor Superfamily Member 5A Precursor | CDS/nonSyn |
| 7.79 | 16 | 39966611 | 666 | + | tnfrsf5a | Tnf Receptor Superfamily Member 5A Precursor | 3'UTR |
| 8.47 | 16 | 39967277 | 149527 | + | tnfrsf5a | Tnf Receptor Superfamily Member 5A Precursor | 3'UTR |
| 8.76 | 16 | 40116804 | 438 | NA | NA | NA | NA |
| 9.06 | 16 | 40117242 | 5021 | - | LOC110492067 | Kelch Protein 21 | CDS/syn |
| 9.04 | 16 | 40122263 | 471 | - | LOC110492067 | Kelch Protein 21 | CDS/syn |
| 9.04 | 16 | 40122734 | 206269 | - | LOC110492067 | Kelch Protein 21 | CDS/syn |
| 8.97 | 16 | 40329003 | 423 | + | LOC110492070 | 45 Kda Calcium-Binding Protein | 3'UTR |
| 8.88 | 16 | 40329426 | 430961 | + | LOC110492070 | 45 Kda Calcium-Binding Protein | 3'UTR |
| 8.88 | 16 | 40760387 | 133719 | + | LOC100136676 | Caspase-9 | CDS/syn |
| 8.87 | 16 | 40894106 | 16043 | + | LOC110491067 | Basement Membrane-Specific Heparan Sulfate<br>Proteoglycan Core Protein | CDS/syn |
| 8.81 | 16 | 40910149 | 15660 | + | LOC110491067 | Basement Membrane-Specific Heparan Sulfate<br>Proteoglycan Core Protein | CDS/nonSyn |
| 8.81 | 16 | 40925809 | 134 | NA | NA | NA | NA |
| 8.82 | 16 | 40925943 | 328 | NA | NA | NA | NA |
| 8.88 | 16 | 40926271 | 36300 | NA | NA | NA | NA |
| 8.88 | 16 | 40962571 | 1603 | + | LOC110492082 | Cdp-Diacylglycerol--Serine O-Phosphatidyltransferase | 3'UTR |
| 8.88 | 16 | 40964174 | 1011 | NA | NA | NA | NA |
| 8.89 | 16 | 40965185 | 134 | NA | NA | NA | NA |
| 8.89 | 16 | 40965319 | 214946 | NA | NA | NA | NA |
| 9.33 | 16 | 41180265 | 15995 | + | LOC110492084 | Membrane-Associated Guanylate Kinase, Ww And<br>PdZ Domain-Containing Protein 3 | CDS/syn |
| 9.37 | 16 | 41196260 | 49825 | - | LOC110492085 | Tyrosine-Protein Phosphatase Non-Receptor Type 12 | CDS/syn |
| 9.82 | 16 | 41246085 | 3112 | + | LOC100136105 | Complement Receptor | CDS/syn |
| 9.82 | 16 | 41249197 | 474 | + | LOC100136105 | Complement Receptor | 3'UTR |
| 9.83 | 16 | 41249671 | 30475 | + | LOC100136105 | Complement Receptor | 3'UTR |
| 9.95 | 16 | 41280146 | 574 | + | c4bp | C4B-Binding Protein Alpha Chain | 3'UTR |
| 9.93 | 16 | 41280720 | 774 | + | c4bp | C4B-Binding Protein Alpha Chain | 3'UTR |
| 10.15 | 16 | 41281494 | 229 | NA | NA | NA | NA |
| 10.29 | 16 | 41281723 | 24001 | NA | NA | NA | NA |
| 10.33 | 16 | 41305724 | 20095 | - | LOC110492088 | Uncharacterized Loc110492088 | NA |
| 10.36 | 16 | 41325819 | 685099 | - | cd34a | Cd34A Molecule | 3'UTR |
| 10.44 | 16 | 42010918 | 5137 | - | slc26a9 | Solute Carrier Family 26 Member 9 | CDS/nonSyn |
| 10.45 | 16 | 42016055 | 176696 | - | slc26a9 | Solute Carrier Family 26 Member 9 | CDS/syn |
| 10.49 | 16 | 42192751 | 41683 | - | LOC110492098 | Cysteine/Serine-Rich Nuclear Protein 2 | CDS/syn |
| 9.58 | 16 | 42234434 | 23274 | + | LOC110492102 | Daz-Associated Protein 2 | 3'UTR |
| 9.68 | 16 | 42257708 | 1026 | - | LOC110492103 | Rac Gtpase-Activating Protein 1 | 3'UTR |
| 9.45 | 16 | 42258734 | 38505 | - | LOC110492103 | Rac Gtpase-Activating Protein 1 | 3'UTR |
| 8.01 | 16 | 42297239 | 2891 | + | LOC110492108 | Citrate Synthase, Mitochondrial | CDS/nonSyn |
| 8.01 | 16 | 42300130 | 5927 | + | LOC110492108 | Citrate Synthase, Mitochondrial | CDS/nonSyn |
| 7.38 | 16 | 42306057 | 101 | + | LOC110492108 | Citrate Synthase, Mitochondrial | 3'UTR |
| 6.57 | 16 | 42306158 | 92 | + | LOC110492108 | Citrate Synthase, Mitochondrial | 3'UTR |
| 6.01 | 16 | 42306250 | 1 | + | LOC110492108 | Citrate Synthase, Mitochondrial | 3'UTR |
| 5.19 | 16 | 42306251 | 60 | + | LOC110492108 | Citrate Synthase, Mitochondrial | 3'UTR |
| 4.54 | 16 | 42306311 | 303 | + | LOC110492108 | Citrate Synthase, Mitochondrial | 3'UTR |
| 3.90 | 16 | 42306614 | 605 | + | LOC110492108 | Citrate Synthase, Mitochondrial | 3'UTR |
| 3.19 | 16 | 42307219 | 57 | + | LOC110492108 | Citrate Synthase, Mitochondrial | 3'UTR |
| 2.53 | 16 | 42307276 |  | + | LOC110492108 | Citrate Synthase, Mitochondrial | 3'UTR |

**Figure 4.**
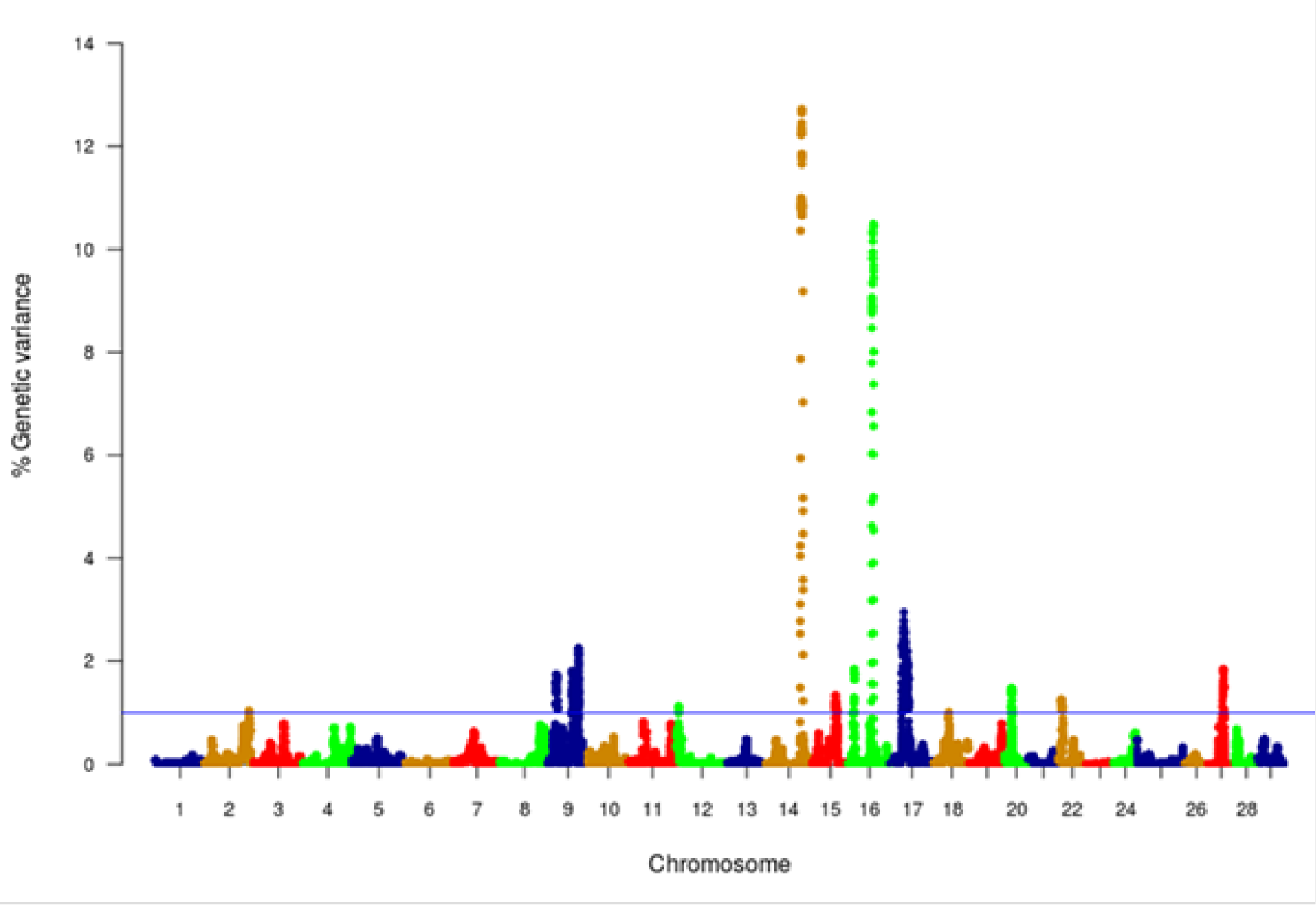
Manhattan plot of GWA analysis performed with WssGBLUP and showing association between SNP genomic sliding windows of 50 SNPs and muscle yield. Chromosomes 14 and 16 showed the highest peaks with genomic loci, explaining together up to 23.2% of the genetic variance. The blue line shows a threshold of 1% of additive genetic variance explained by SNPs.

With 46 SNPs clustered into 23 annotated genes, chromosome 14 had the most significant QTL windows explaining up to 12.71% of the genetic variance in muscle yield (Table 3 and Figure 4). At least four genes can be inferred to be involved in cell differentiation/proliferation and regulation of gene expression based on their RefSeq annotation. The list included fibroblast growth factor-binding protein-1(FGFBP1) which had a single nonsynonymous SNP found in a window that explained 12.24% of the additive genetic variance. FGFBP1 plays an essential role in cell proliferation and differentiation by binding to fibroblast growth factors. The FGFBP1 expression increases during development and decreases before neuromuscular junction degeneration during aging [33]. The list of genes on chromosome 14 also includes inositol polyphosphate 5-phosphatase (OCRL-1). OCRL is involved in terminating the PI3K signaling and thus plays an important role in modulating effects of growth factors and insulin stimulation in cell proliferation and survival [34]. Prominin-1-A gene (PROM1) that encodes for a transmembrane glycoprotein had 2 SNPs. PROM1, often used as adult stem cell marker, plays a role in maintaining stem cell properties by suppressing differentiation [35]. Another gene on chromosome 14 was farnesyltransferase/geranylgeranyltransferase type-1 subunit alpha (FNTA) which had a SNP explaining 12.36% of the variance. FNTA may positively regulate neuromuscular junction development [36].

In addition, chromosome 14 had three genes involved in the cell cycle regulation. The first gene is MCTS1 re-initiation and release factor that had two SNPs in a window explaining 12.65% of the additive genetic variance. MCTS1 is anti-oncogene that decreases cell doubling time by shortening the G1 and G1/S transit time [37]. The second cell cycle control gene was cyclin-A2 which promotes transition through G1/S and G2/M and can block muscle-specific gene expression during muscle differentiation [38]. The third gene was glutathione S-transferase P (GSTP1).

Although involved in numerous biological functions, GSTP1 negatively regulates CDK5 activity via p25/p35 translocation which diminishes neurodegeneration [39].

Chromosome 14 also had SNPs in genes playing important mitochondrial functions. There were 4 SNPs in the gene encoded for the electron transfer flavoprotein dehydrogenase (ETFDH) which is an important enzyme in the mitochondrial electron transport chain. Mutations in ETFDH are associated with myopathies[40]. Another mitochondrial-relevant gene was peptidylprolyl isomerase D (PPID). Mutations in PPID are associated with muscular dystrophy in human [41].

Few other genes included in the QTL region on chromosome 14 are important for maintenance of the muscle functions. Of them is the chloride intracellular channel protein 2 (CLIC2) which modulates the activity of ryanodine receptor 2 (RYR2) and inhibits calcium influx, and therefore is involved in regulating muscle contraction [35]. Five SNPs were in the lysosomal-associated membrane protein 2 gene (LAMP2). LAMP2 mutations were reported in patients with cardioskeletal myopathies [36]. Two SNPs were located in the UPF3B gene, a regulator of nonsense-mediated mRNA decay (NMD). NMD inhibition was observed in patients with muscular dystrophy [37]. Three SNPs were observed in the septin-6 gene. Mutations of septin-9 (another gene family member) is genetically linked to muscle atrophy [38]. Two SNPs were identified in the tenomodulin gene which showed downregulation in an animal muscle atrophy model [39].

Chromosome 16 ranked second in having the most significant QTL windows with 49 SNPs clustered into 16 annotated genes (Table 4 and Figure 4). The gene within the most significant SNP window to additive genetic variance was the cysteine/serine-rich nuclear protein 2 (CSRNP2). CSRNP2 has DNA binding transcription factor/activation activity. Deletion of CSRNP1/2/3 three gene family members resulted in mice neonatal lethality [42]. Another gene within the same SNP window was solute carrier family 26 member 9 (Slc26a9). Little is known about the function of Slc26a9 in muscle, it serves as anion exchanger mediating chloride, sulfate and oxalate transport and chloride/bicarbonate exchange [43]. A single SNP was observed in the stem cell marker CD34a gene. Cd34(-/-) mice showed a defect in muscle regeneration caused by acute or chronic muscle injury [44].

Several genes were involved in cell signaling/receptor activity. Five SNPs were predicted in 2 genes of the immune-related complement activation pathway, these are the complement receptor and C4b-binding protein alpha chain. Recent studies indicated that the complement is activated as a response of skeletal muscle injury and plays a key role during muscle regeneration[45]. A single SNP was identified in the tyrosine-protein phosphatase non-receptor type 12 (PTPN12) which dephosphorylates a wide-range of proteins, and thus regulates several cellular signaling cascades such as ERBB2 and PTK2B/PYK2 [46]. This group of genes also includes the membrane-associated guanylate kinase, WW and PDZ domain-containing protein 3 (MAGI3), which is involved in the regulation of various cell signaling processes including the AKT1, TGFA, ERK and JNK signaling cascades [47]. Two SNPs were in the basement membrane-specific heparan sulfate proteoglycan core protein (HSPG2). A mouse model deficient in this gene showed muscle hypertrophy through reduced myostatin expression suggesting a role in maintaining fast muscle mass and fiber composition [48]. Five SNPs were in the TNF receptor superfamily member 5A gene. Recently, some proinflammatory cytokines belonging to TNF superfamily have been recognized as an important regulator of skeletal muscle mass [49].

Chromosome 16 also had a single SNP in the DAZ-associated protein 2 (DAZAP2). Not much is known about the DAZAP2 function in muscle, however, DAZAP2 interacts with the transforming growth factor-beta signaling molecule SARA (Smad anchor for receptor activation), eukaryotic initiation factor 4G, and an E3 ubiquitinase [47]. Another gene in the list was Rac GTPase-activating protein 1 (RACGAP1) that harbored 2 SNPs explaining up to 9.658% of the genetic variance. RACGAP1 regulates cytokinesis and cell differentiation [50]. A single SNP existed in caspase-9 which has an important non-apoptotic role in muscle differentiation [51]. Three SNPs were located in the kelch protein 21. Several Kelch family members play important roles in skeletal muscle development by regulating the cell proliferation and/or differentiation [52].

An important gene affecting muscle function which is also located within the QTL region on chromosome 16 is the citrate synthase (CS), which is used as a marker for human mitochondrial functions. Ten SNPs explaining up to 8% of the genetic variance were located in the CS gene.

GWA studies in fish to identify QTL affecting muscle yield and quality are still in its infancy. Previous GWA analysis using a 57K genomic SNP chip on the same fish population identified two windows that explained 1.5% and 1.0% of the additive genetic variance for muscle yield and 1.2% and 1.1% for muscle weight. Interestingly, the windows are located on chromosome 9, which showed some association with muscle yield in the current study; however, none of the SNPs were annotated to the same genes. No major QTLs were identified in the previous study. This large difference in the outcomes of the two studies was somewhat unexpected. However, it may be explained by lower marker density within or near genes in the 57K chip [10] and by ascertainment bias, because the transcribed SNPs used in this study were discovered in the phenotyped fish and hence are expected to be more polymorphic and informative for GWA analysis in this population. Additionally, in this study, sliding windows of 50 SNP were used contrasting with 20 non-sliding windows in the previous study. Difference in window size slightly contributed to the increased proportion of variance (data not shown). By using SNP windows, it is assumed that those DNA blocks may be inherited together, which may not always be the case for all assumed windows. In common carp, genetic linkage mapping identified QTLs with large effects for muscle fiber cross-section area (21.9%) and muscle fiber density (18.9%) [53]. Genome-wide significant QTL affecting growth and muscle related traits were identified in Atlantic salmon [54]. The latter two studies, together with our study, indicate existence of large-effect QTLs affecting muscle yield in aquaculture species. However, the QTLs identified in this study might be population specific and thus, need to be tested in other populations.

#### Citrate synthetase activity correlation with muscle yield

A SNP window on chromosomes 16 explaining up to 8.01% of the genetic variance in muscle yield contained ten SNPs of the CS gene. Two of the SNPs were nonsynonymous mutations. To investigate the potential effect of these SNPs, we measured the CS activity in 100 fish from the 2012 year-class. The samples included 38 fish from 5 high-ranked and 5 low-ranked families for muscle yield and 62 randomly selected fish. CS had 1.43-fold increase in the high-ranked fish compared to the low ranked ones (figure 5). The regression coefficient R^2^value between the muscle yield and CS activity was 0.092 (p-value 0.002). These results indicate an important role of mitochondrial functions to muscle growth. Mitochondria are at the center of age-related sarcopenia that is characterized by decline in human muscle mass. Skeletal muscle CS decreases with aging in humans [55].

**Figure 5.**
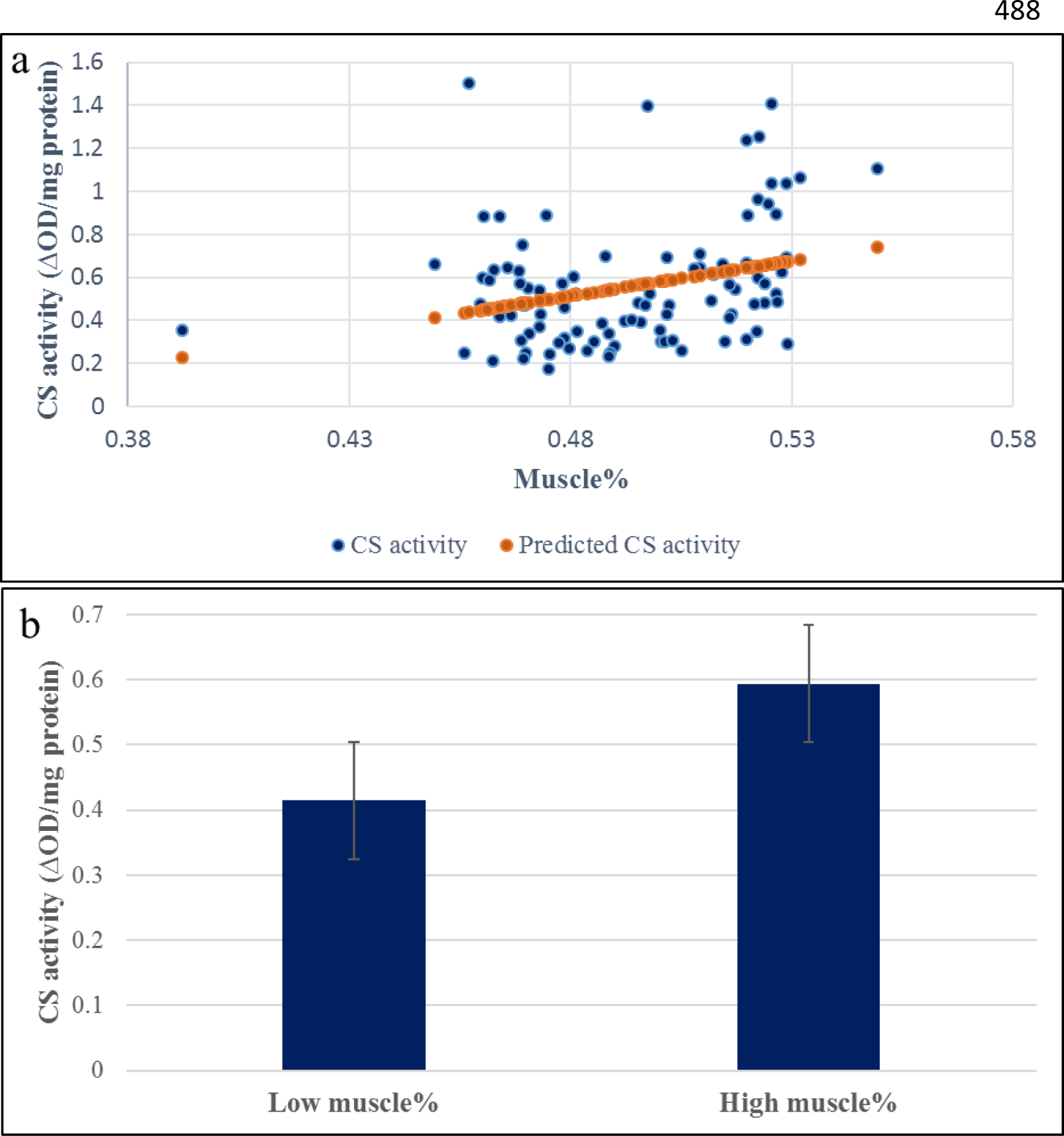
Correlation coefficient between muscle yield and CS activity in 96 samples. A) The regression coefficient R2 value between the muscle yield and CS activity was 0.092 (p-value 0.002). b) CS had 1.43-fold increase in the high-ranked fish compared to the low ranked ones.

### CONCLUSION

This study provides a 50K transcribed gene SNP-chip based on RNA-Seq data from fish families showing genetic diversity for six aquaculture production traits in the USDA/NCCCWA growth- and disease-selected genetic lines. The chip was tested for GWA analysis, which led to identification of large-effect QTL for muscle yield in that population. Other muscle quality traits are currently under investigation. Collectively, these studies will allow the use of SNP markers to estimate breeding values for muscle yield and quality traits that are economically important traits for aquatic food producers, processors, and consumers. Current and future selection at the NCCCWA will select for improved fillet yield. Genetic markers are desirable for these traits because genetic improvement is limited by the inability to measure fillet yield traits directly on broodstock due to lethal sampling. Hence the accuracy and efficiency of selective breeding can be improved by taking advantage of the genomic information, even though limited phenotyping is available for this economically-important trait.

## AUTHOR CONTRIBUTIONS

Conceived and designed the experiments: MS, TL, BK. Performed the experiments: RA, MS, TL, BK. Analyzed the data: RA, AA, DL, GG, YP, BK, MS. Wrote the paper: MS. All authors reviewed and approved the publication.

## CONFLICT OF INTEREST STATEMENT

The authors declare that they have no competing interests.

## ACKNOWLEDGMENTS

This study was supported by a competitive grant No. 2014-67015-21602 from the United States Department of Agriculture, National Institute of Food and Agriculture (MS). R.A.T trainee’s projects are supported by Grant Number T32HL072757 from the National Heart, Lung, and Blood Institute. The content is solely the responsibility of the authors and does not necessarily represent the official views of any of the funding agents.

